# Expression of mRNAs Encoding Hypothalamic Small Proteins, Neurosecretory Protein GL and Neurosecretory Protein GM, in the Japanese Quail, *Coturnix japonica*

**DOI:** 10.1101/2023.09.01.552892

**Authors:** Masaki Kato, Eiko Iwakoshi-Ukena, Yuki Narimatsu, Megumi Furumitsu, Kazuyoshi Ukena

## Abstract

Neurosecretory protein GL (NPGL) and neurosecretory protein GM (NPGM) are novel neuropeptides that have been discovered in the hypothalamic infundibulum of chickens. NPGL and NPGM play important roles in lipid metabolism in juvenile chickens. The physiological functions of NPGL and NPGM in sexually mature birds remain unknown. The Japanese quail (*Coturnix japonica*) seems to be an appropriate model for analyzing NPGL and NPGM during sexual maturity. However, studies on NPGL or NPGM have yet to be reported in the Japanese quail. In the present study, we identified cDNAs encoding precursor proteins of NPGL and NPGM in the quail hypothalamus. In situ hybridization revealed that *NPGL* mRNA-expressing cells in the hypothalamus were localized in the infundibular nucleus and median eminence, and *NPGM* mRNA-expressing cells were only found in the mammillary nucleus. Immunohistochemistry revealed that NPGM-like immunoreactive cells were distributed in the mammillary nucleus, whereas NPGL-like immunoreactive cells were not detected in the hypothalamus. Real-time PCR analysis indicated that the expression of *NPGL* mRNA was higher in the hypothalamus of females than in males, and *NPGM* mRNA expression showed no sex differences. *NPGL* and *NPGM* mRNA expression in males was upregulated after 24 h of food deprivation. In females, only *NPGM* mRNA expression was increased by fasting. These results suggest that the physiological functions of NPGL and NPGM are different in quail, and these factors are involved in sex differences in energy metabolism.

## INTRODUCTION

Energy homeostasis is maintained by a balance between food intake, energy expenditure, and energy storage (Morton et al., 2014). In mammals, food intake and energy metabolism are regulated by neuropeptides and/or hormones secreted from the brain and peripheral tissues (Schwartz et al., 2000). Neuropeptides involved in feeding behavior and energy metabolism are secreted by neurons in the hypothalamus (Huang et al., 2021). Similar to mammals, the hypothalamus in birds is also an energy metabolic center. However, the regulatory mechanisms of feeding behavior differ between birds and mammals (Tachibana and Tsutsui, 2016). For instance, orexin and melanin-concentrating hormone (MCH) are orexigenic factors in mammals (Garthwaite, 1985; Sakurai et al., 1998), whereas these neuropeptides do not affect feeding behavior in neonatal chicks (Furuse et al., 1999; Ando et al., 2000). Growth hormone-releasing hormone (GHRH) stimulates the release of growth hormone (GH) in the pituitary gland and promotes feeding behavior in mammals (Vaccarino et al., 1985; Sherwood et al., 2000). However, in neonatal chicks, GHRH suppresses feeding behavior (Tachibana et al., 2015). Thus, the mechanism of appetite control in birds is unique and not fully understood. Therefore, it is likely that unknown factors are involved in these processes.

We discovered a novel cDNA encoding the precursor of a small secretory protein in the chicken hypothalamus (Ukena et al., 2014). The novel secretory protein was named neurosecretory protein GL (NPGL) because the predicted C-terminal amino acid of the small protein was Gly-Leu-NH2. The precursor gene for *NPGL* is highly conserved in vertebrates (Matsuura et al., 2017; Ukena, 2018; Huang et al., 2022).

*NPGL* mRNA-expressing cells and NPGL-like immunoreactive cells are localized in the mammillary nucleus (MM) and infundibular nucleus (IN) of the chicken hypothalamus (Ukena et al., 2014; Shikano et al., 2018b). In mice, *NPGL* mRNA-expressing cells are localized in the lateroposterior part of the arcuate nucleus (ArcLP) of the hypothalamus (Matsuura et al., 2017). In GIFT tilapia (*Oreochromis niloticus*), *NPGL* mRNA-containing cells are localized in the anterior periventricular pretectal nucleus of the anterior preoptic area and lateral tuberal nucleus of the hypothalamus (Huang et al., 2022). We previously showed that NPGL induces food intake and fat accumulation in rats, mice, and chicks (Iwakoshi-Ukena et al., 2017; Matsuura et al., 2017; Shikano et al., 2019; Narimatsu et al., 2022a, b). These results suggest that NPGL is a novel regulator of energy metabolism.

*NPGL* gene has another paralogous gene, neurosecretory protein GM (*NPGM*), in chickens. *NPGM* and *NPGL* genes are conserved in vertebrates, including chickens, rats, and humans (Ukena et al., 2014). The predicted C-terminal amino acid of the mature NPGM peptide was Gly-Met-NH2 (Shikano et al., 2018a). In chicks, *NPGM* mRNA-expressing cells and NPGM-like immunoreactive cells are localized in the MM and IN. Moreover, it is possible that NPGM and NPGL are produced in the same neuron of the MM and IN (Shikano et al., 2018a). In addition to histological analyses, we previously reported that NPGM could play significant roles in fat accumulation and stress response in chicks (Kato et al., 2021, 2022). In mice, intracerebroventricular (i.c.v.) administration of NPGM promotes feeding behavior during dark periods (Martinez et al., 2023). These results suggest that NPGM is related to energy metabolism as well as NPGL in higher vertebrates.

Although physiological analyses of NPGL and NPGM in birds have been performed by chronic i.c.v. administration using juvenile chicks (Shikano et al., 2019; Kato et al., 2021), this method has not been established in adult chickens (Lamberigts et al., 2022). Therefore, the biological functions of NPGL and NPGM in birds after sexual maturity remain unknown. The Japanese quail (*Coturnix japonica*) is not only an important livestock animal, but also an appropriate animal model for the study of neuroendocrinology (Yoshimura et al., 2003; Tsutsui, 2016). The advantages of the Japanese quail as a model animal are as follows: they are small (∼17 cm), mature quickly (6–8 weeks), and can be kept in large numbers at the laboratory (Baer et al., 2015). Therefore, the Japanese quail seems to be suited for the elucidation of the physiological functions of NPGL and NPGM in energy metabolism in adult birds.

However, the presence of NPGL and NPGM in quails has not yet been reported. In the present study, we identified cDNAs encoding precursor proteins of NPGL and NPGM in the hypothalamus of quails. Second, morphological analyses using in situ hybridization and immunohistochemistry were employed for the localization of NPGL and NPGM in the hypothalamus. Third, *NPGL* and *NPGM* mRNA expression levels were compared in males and females using real-time PCR. Finally, the expression of *NPGL* and *NPGM* mRNA was examined in males and females after fasting. In the gene expression analysis by real-time PCR, previously known feeding-related factors were also analyzed to examine their relationship with novel factors, NPGL and NPGM, in this study.

## MATERIALS AND METHODS

### Animals

Fertilized eggs of the Japanese quail (*Coturnix japonica*) were purchased from a commercial company (Motoki Corporation, Saitama, Japan) and hatched in the laboratory. Post-hatched male and female quails were housed in a windowless room at 25°C under long-day conditions (16 h light and 8 h dark) with ad libitum access to water and a commercial diet (Dream Co., Ltd., Aichi, Japan). The experimental protocols were in accordance with the Guide for the Care and Use of Laboratory Animals prepared by Hiroshima University (Higashi-Hiroshima, Japan), and the procedures were approved by the Institutional Animal Care and Use Committee of Hiroshima University (permit number C22-27).

### Sequencing of the cDNA of *NPGL* and *NPGM* in Japanese quail

We searched the sequences of quail *NPGL* and *NPGM* using a BLAST search based on the chicken sequences (chicken *NPGL*; NM_001389496.2, chicken *NPGM*; XM_429770.2). The predicted sequence of quail *NPGL* was registered as *Coturnix japonica* protein FAM237A-like (LOC107314652) for mRNA (XM_015864121.2). A similar sequence of quail *NPGM* was registered as *Coturnix japonica* uncharacterized LOC116653024 (LOC116653024) for ncRNA (XR_004306394.1). Based on these preliminary data, primers for sequencing cDNAs encoding the open reading frame were designed from the predicted nucleotide sequences. The primer sets used are listed in Table 1.

**Table 1.**
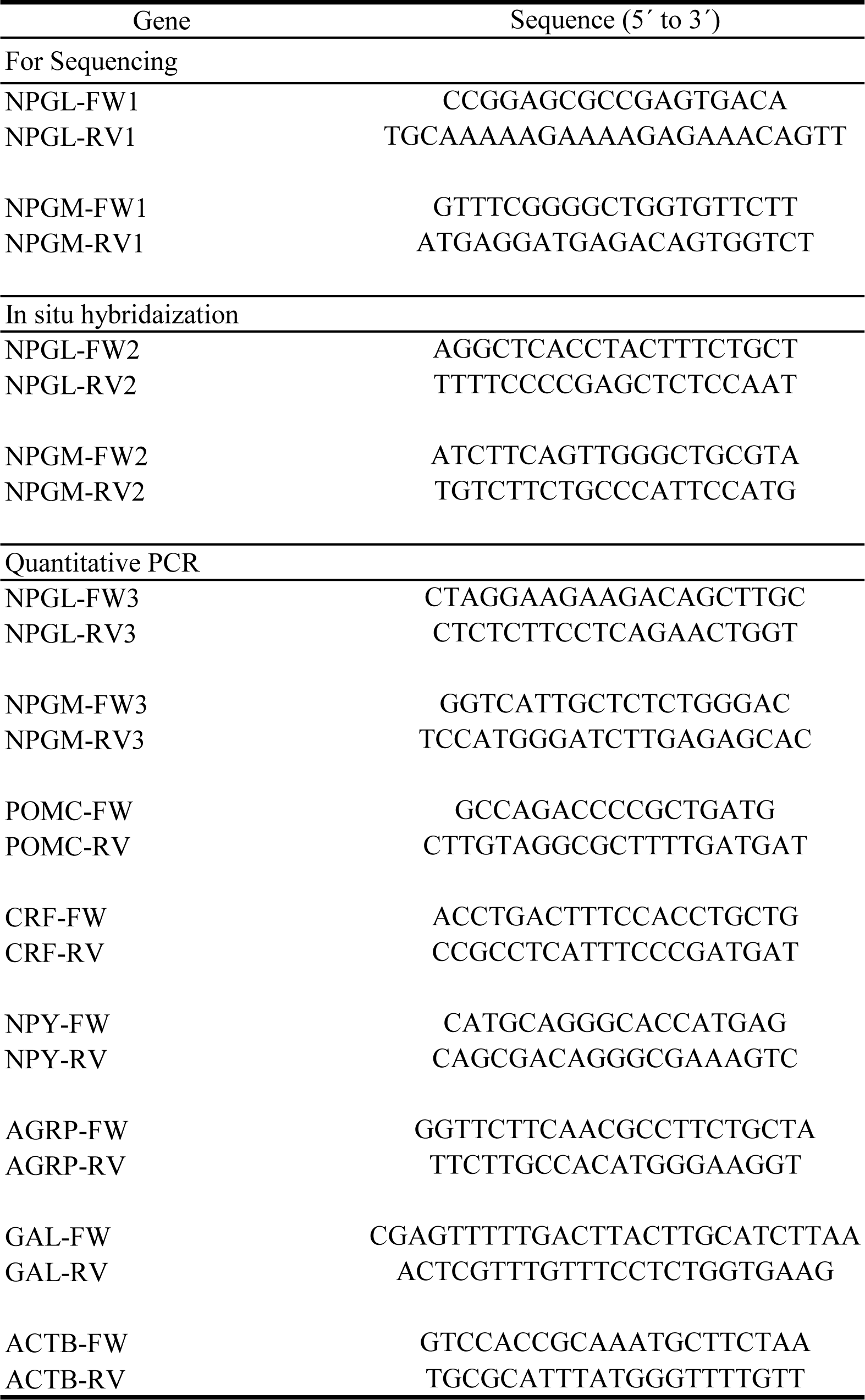
The list of primer sets used in this study.

The hypothalamic infundibulum was dissected and snap-frozen in liquid nitrogen for further RNA processing from five day-old male or female quails. Total RNA from the hypothalamic infundibulum was isolated using TRIzol reagent (Thermo Fisher Scientific, Waltham, MA) according to the manufacturer’s instructions. First-strand cDNA was synthesized from total RNA using the PrimeScript RT Reagent Kit with gDNA Eraser (Takara Bio, Shiga, Japan). PCR amplifications were performed using the KOD One PCR Master Mix (TOYOBO, Osaka, Japan). The PCR products were purified using Nucleospin Gel and PCR Clean-up (Takara Bio). Nucleotide sequence analysis was outsourced from a commercial sequencing service (Eurofins, Tokyo, Japan). Protein sequence alignments were performed using Snap Gene Viewer (https://www.snapgene.com/).

### In situ hybridization for *NPGL* and *NPGM*

Primers for antisense and sense RNA probes were designed based on the sequences of quail *NPGL* and *NPGM* (Table 1). Digoxigenin-labeled antisense and sense RNA probes were produced from cloned PCR fragments using a digoxigenin RNA labeling kit (SP6/T7; Roche Diagnostics, Basal, Switzerland). Labeling was performed as previously described (Ukena et al., 2008, 2010).

In situ hybridization was performed in a manner similar to that described previously (Ukena et al., 1999). Male quails (nine week-old) were transcardially perfused with 0.9% saline, followed by 4% paraformaldehyde. The brains were postfixed overnight and then put in sucrose solution (30% sucrose in 10 mM phosphate buffer) at 4°C until they sank. The brains were cut at 10 µm thickness with a cryostat at −20°C. The frozen sections were rehydrated in PBS and treated with 0.2 N HCl for 20 min, followed by 1 μg/ml proteinase K at 37°C for 10 min. After fixation in 4% paraformaldehyde for 5 min, the sections were maintained in 40% deionized formamide in 4× SSC (1× SSC = 150 mM NaCl, 15 mM sodium citrate, pH 7.0) for 30 min. Hybridization was performed overnight at 50°C with 50 ng/ml digoxigenin- oligonucleotide probe mixture dissolved in the hybridization medium (10 mM Tris-HCl (pH 7.4), 1 mM EDTA, 0.6 M NaCl, 10% dextran sulfate, 1 × Denhardt’s solution, 250 μg/ml yeast transfer RNA, 125 μg/ml salmon sperm DNA, and 40% deionized formamide). After hybridization, the sections were washed six times with 50% formamide-2 × SSC at 55°C for 10 min each time. After blocking with a blocking solution (1% BSA in PBS containing 0.3% Triton X-100) for 30 min, the sections were incubated with alkaline phosphatase-labeled anti-digoxigenin antibody (1:1000 dilution in the blocking solution) for 1 h. After incubation, the sections were washed three times with 0.075% Brij 35 in PBS for 10 min each time. Single staining was performed with nitro blue tetrazolium/5-bromo-4-chloro-3-indolyl phosphate (NBT/BCIP; 1:50 dilution in alkaline buffer; Roche Diagnostics). The cells were then stained overnight at room temperature.

### Immunohistochemistry

To detect NPGL and NPGM in the quail brain, we used antisera against chicken NPGL and NPGM from previous studies (Shikano et al., 2018a, b). Antisera against NPGL and NPGM were produced in guinea pigs and rabbits, respectively. The antiserum of chicken NPGL was confirmed to specifically recognize synthetic quail NPGL and chicken NPGL by competitive enzyme-linked immunosorbent assay (ELISA). The IC50 values (concentrations yielding 50% displacement) of chicken NPGL antibody in the competitive ELISA were estimated as follows; 16.1 pmol for quail NPGL, 8.4 pmol for chicken NPGL, and more than 1000 pmol for chicken/quail NPGM. The specificity of chicken NPGM antibodies has been analyzed in previous reports (Shikano et al., 2018a). The IC50 values of chicken NPGM antibody were estimated as follows; 94.8 pmol for chicken/quail NPGM, and more than 1000 pmol for chicken and quail NPGL.

The brains of nine week-old male quail were cut at 10 µm thickness with a cryostat at −20°C. Frozen sections were incubated with 0.3% hydrogen peroxide (H2O2) in absolute methanol for 30 min. After blocking with blocking solution (1% BSA in PBS containing 0.3% Triton X-100) for 1 h, the sections were incubated with guinea pig anti- NPGL antibody (1:500 dilution in blocking solution) or rabbit anti-NPGM antibody (1:1000 dilution in blocking solution) overnight at 4°C. Biotinylated goat anti-guinea pig IgG (1:1000 dilution in blocking solution, Vector Laboratories, Burlingame, CA) for the detection of NPGL and biotinylated goat anti-rabbit IgG (1:1000 dilution in blocking solution, Vector Laboratories) for the detection of NPGM were used as secondary antibodies. Immunoreactivity was detected with an ABC kit (VECTASTATIN Elite Kit; Vector Laboratories), followed by a diaminobenzidine (DAB) reaction. In control experiments, pre-adsorption was performed as a control using the antigen (10 µg/mL NPGL or NPGM).

### Comparison of gene expression levels between males and females

Male quails (n = 5) and female quails (n = 5) at six weeks of age were used to compare gene expression levels between males and females. Body mass and blood glucose levels were measured before euthanasia. Blood samples were collected from the ulnar veins and analyzed using a GLUCOCARD G+ meter (Arkray, Kyoto, Japan). The hypothalamic infundibulum was collected, immediately snap-frozen in liquid nitrogen, and stored at −80°C for real-time PCR analysis.

### 24 hours Food Deprivation Experiment

Male quails (n = 5) and female quails (n = 5) at six weeks of age were fasted for 24 h. The quails had access to water ad libitum. Control quails (male and female, n = 5 each) had ad libitum access to food and water. Before fasting, body mass and blood glucose levels were measured. Blood samples were collected from the ulnar veins and analyzed using a GLUCOCARD G+ meter (Arkray). After fasting, body mass and blood glucose levels were measured again. The quails were euthanized, and the hypothalamic infundibulum was collected, immediately snap-frozen in liquid nitrogen, and stored at −80°C for real-time PCR analysis.

### Quantitative RT-PCR

RNA was extracted from the hypothalamic infundibulum using Nucleospin RNA Plus XS (Takara Bio). PCR amplifications were performed using THUNDERBIRD SYBR qPCR Mix (TOYOBO): 95°C for 20 s, followed by 40 cycles at 95°C for 3 s and 60°C for 30 s using a real-time thermal cycler (CFX Connect; BioRad, Hercules, CA).

Primer sets for *NPGL* and *NPGM* for quantitative PCR were designed based on the results of cDNA sequencing. In addition, we chose the anorexigenic factors, pro- opiomelanocortin (*POMC*) and corticotropin-releasing factor (*CRF*), and orexigenic factors, neuropeptide Y (*NPY*), agouti-related peptide (*AGRP*), and galanin (*GAL*) (Lear et al., 2017; McConn et al., 2019). The primer sets used are listed in Table 1. The relative quantification of each expression was determined by the 2^−ΔΔ*Ct*^ method using β-actin (*ACTB*) as an internal control.

### Statistical Analysis

Data were analyzed using Student’s t-test for mRNA expression, body weight, and blood glucose levels. The significance level was set at *p* < 0.05. All results are expressed as the mean ± SEM.

## RESULTS

### Identification of cDNAs encoding *NPGL* and *NPGM* in the quail

Sequence analyses showed that *NPGL* and *NPGM* cDNA encoded 185 amino acids (555 bp) and 132 amino acids (396 bp) for the open reading frame, respectively. The sequences of the precursor proteins of NPGL and NPGM are shown in Figure 1. The N-terminal signal sequences of the precursor proteins were 32 and 24 amino acid signal peptides, respectively. The mature NPGL and NPGM proteins were predicted to have 80 and 83 amino acid residues, respectively, because their C-terminal amino acid sequences are Gly-Leu-NH2 and Gly-Met-NH2. The precursor proteins possessed a Gly residue, which is an amidation donor, and an Arg-Arg motif, which consists of a potential proteolytic processing site, at the C-terminal. Mature NPGL and NPGM contain two Cys residues. The position of the two Cys residues is also conserved in other vertebrates, suggesting intramolecular disulfide bond formation.

**Fig. 1.**
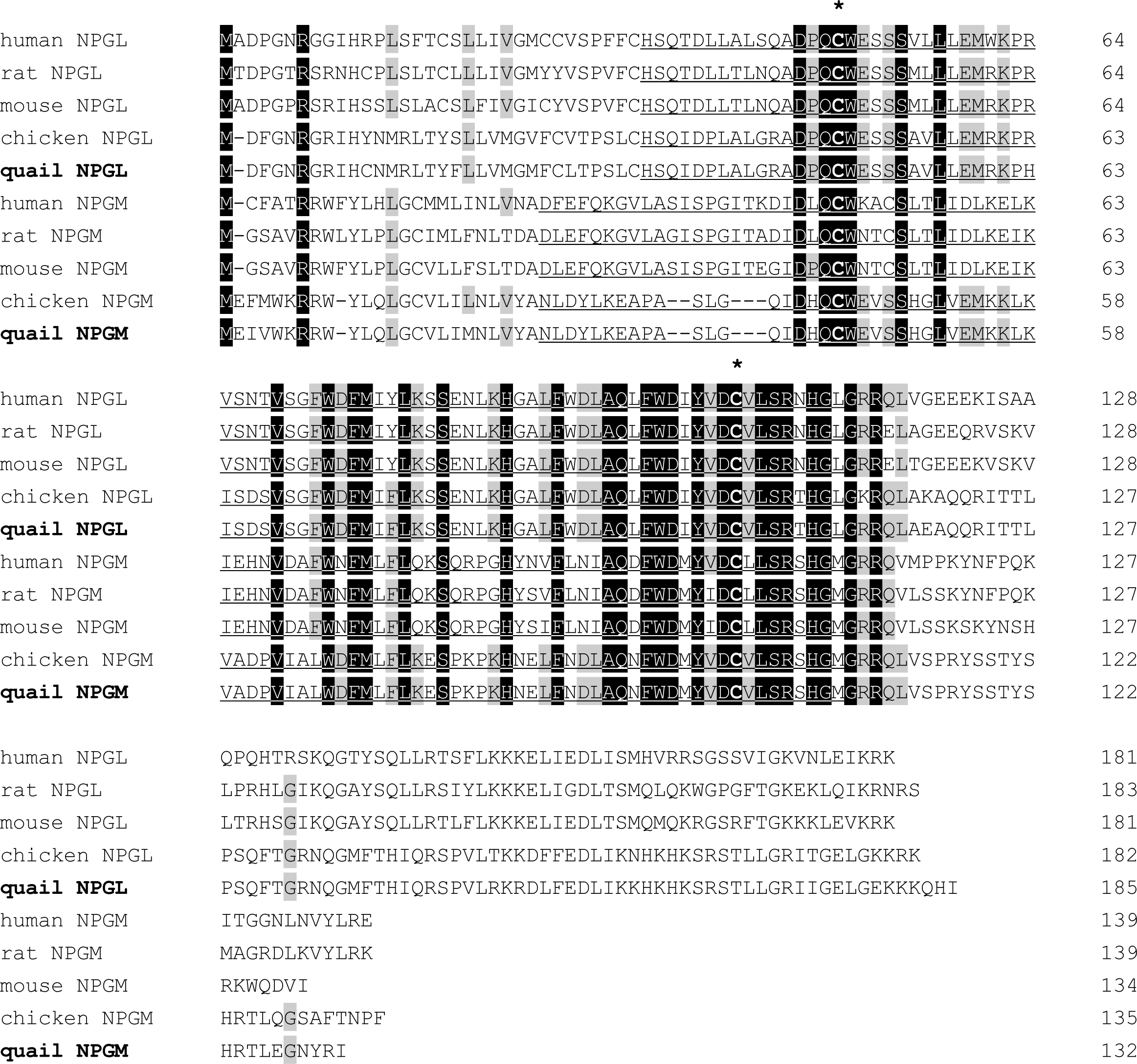
Amino acid sequence alignment of neurosecretory protein GL (NPGL) and neurosecretory protein GM (NPGM) precursor proteins deduced from human, rat, mouse, chicken, and quail cDNA sequences. Black and gray boxes highlight fully conserved and highly conserved amino acids, respectively. Gaps, indicated by hyphens, were inserted to optimize the sequence alignment. The predicted mature small proteins are underlined. The conserved Cys (C) residues are indicated by asterisks.

The similarities of precursor and mature proteins between NPGL and NPGM were 32% and 54%, respectively, in quail. The mature protein of quail NPGL shares high amino acid sequence identity (99%) with chicken NPGL, with only one amino acid substitution. The amino acid sequence similarity of mature NPGL between quail and mammals (humans, rats, and mice) is 84%. The amino acid sequence of mature NPGM between quail and chicken is identical (100%). The amino acid sequence similarity of mature NPGM between quail and humans is 54%, and that of NPGM between quail and rodents (rats and mice) is 52%.

### Localization of *NPGL* and *NPGM* mRNA-expressing cells and immunoreactive cells

*NPGL* and *NPGM* mRNA-expressing cells are localized in the hypothalamic infundibulum. The location of the hypothalamic infundibulum is shown in the coronal brain illustration and mRNA-expressing cells are shown by dots (Fig. 2A, G). In situ hybridization showed that *NPGL* mRNA-expressing cells were localized in the IN and the median eminence (ME) (Fig. 2B, C, D), and *NPGM* mRNA-expressing cells were only localized in the MM (Fig. 2H, I, J). The specificity of the antisense probe was confirmed by its reaction with the sense probe (Fig. 2E, F, K, L).

**Fig. 2.**
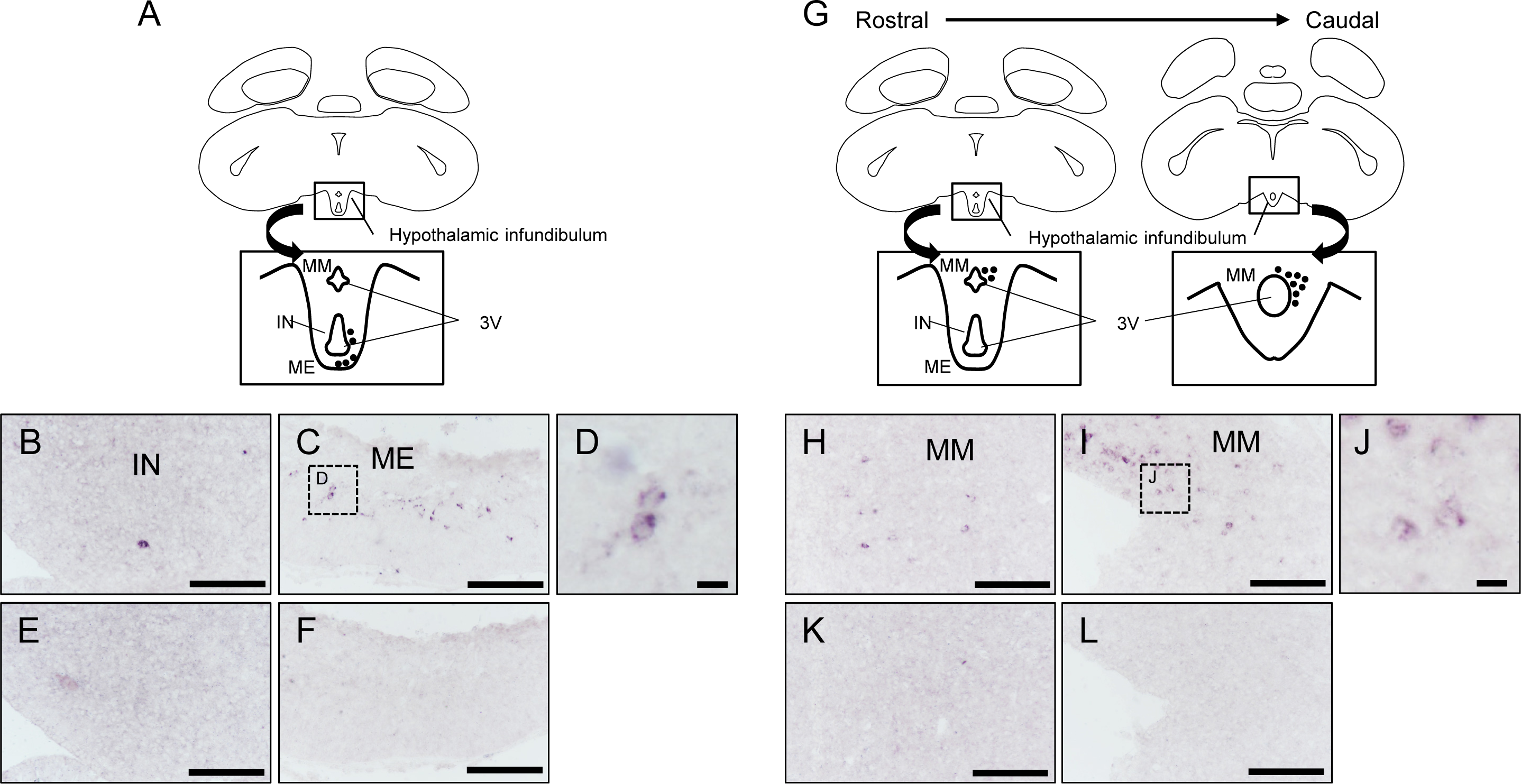
Localization of *NPGL* and *NPGM* mRNA in the hypothalamic infundibulum. (A, G) The location of the hypothalamic infundibulum is shown in the coronal brain illustration. Cells expressing the *NPGL* and *NPGM* precursor mRNA in the medial mammillary nucleus (MM), the infundibular nucleus (IN), and the median eminence (ME) within the hypothalamic infundibulum are represented by dots in the illustration. (B, C) *NPGL* mRNA-expressing cells were localized in the IN and ME. (D) Magnified image of cell bodies of the *NPGL* mRNA-expressing cell in the ME. (E, F) No signals were detected by the sense probe. (H, I) *NPGM*-expressing cells were localized in the MM. (J) Magnified image of cell bodies of the *NPGM* mRNA-expressing cell in the MM. (K, L) No signals were detected by the sense probe. The third ventricle is denoted by 3V. Scale bars = 100 μm in (B, C, E, F, H, I, K, L). Scale bars = 10 μm in (D, J).

The localization and distribution of immunoreactive cells and fibers are shown by dots and dotted lines, respectively (Fig. 3A, F). NPGL-like immunoreactive cells and fibers were not detected in the IN and ME (Fig. 3B, C). However, NPGM-like immunoreactive cells and neural fibers were detected in the MM (Fig. 3G, I, J) and outer layer of the ME (Fig. 3H), respectively. The antibody adsorption test confirmed the disappearance of the immunoreaction (Fig. 3D, E, K, L, M).

**Fig. 3.**
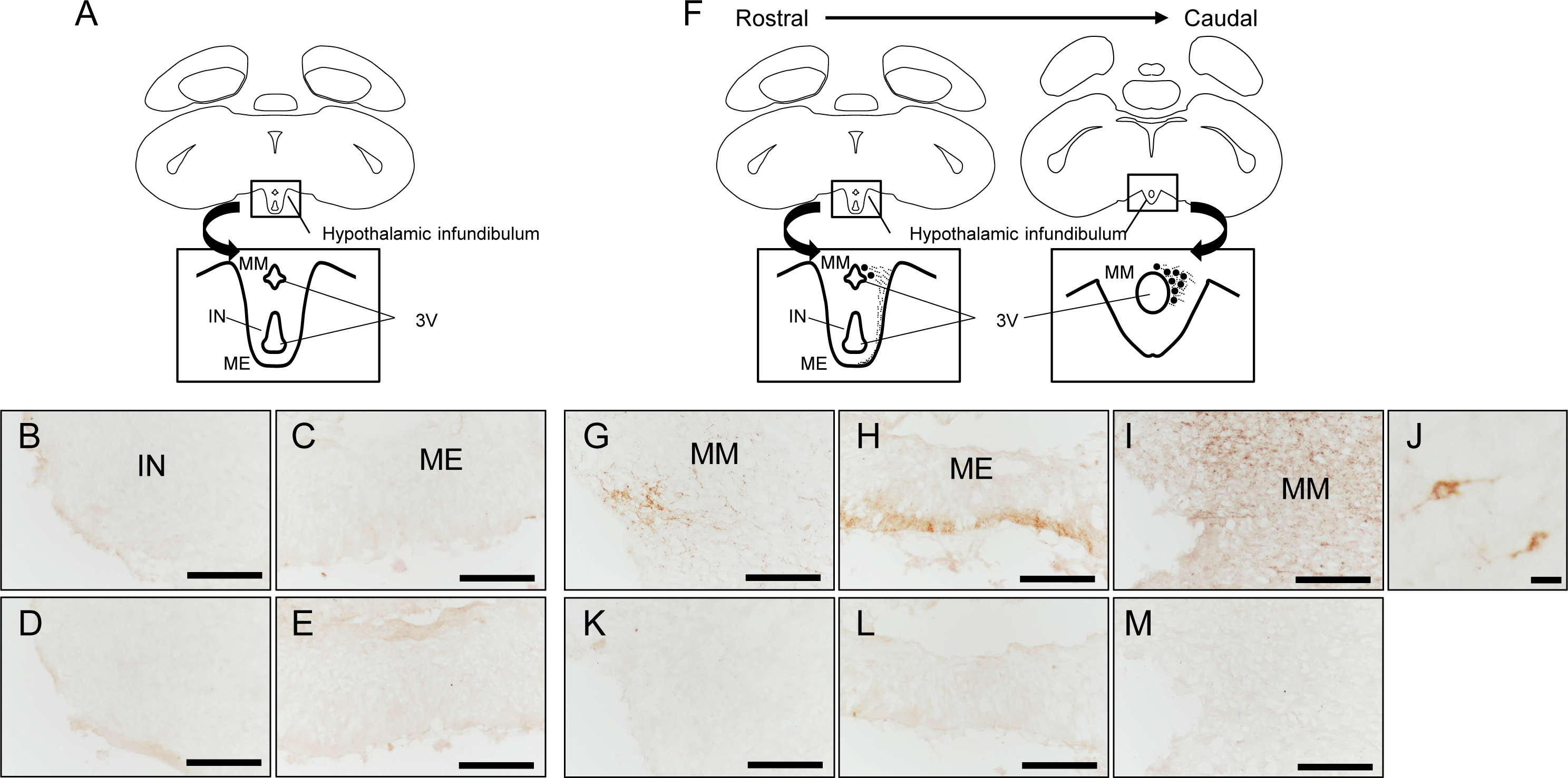
Localization of NPGL and NPGM-like immunoreactive cells in the hypothalamic infundibulum. (A, F) The location of the hypothalamic infundibulum is shown in the coronal brain illustration. NPGM-like immunoreactive cells in the medial mammillary nucleus (MM) within the hypothalamic infundibulum are represented by dots in the illustration. The dotted line is nerve fibers. (B, C) NPGL-like immunoreactive cells in the infundibular nucleus (IN) and the median eminence (ME) were not obtained from nine weeks old male quail. (D, E) No signals were detected by the antibody adsorption test. (G, H, I) NPGM-like immunoreactive cells and nerve fibers in the MM and ME were obtained. (J) Magnified image of the NPGM-like immunoreactive cell body in the MM. (K, L, M) No signals were detected by the antibody adsorption test. The third ventricle is denoted by 3V. Scale bars = 100 μm in (B–I, K–L). Scale bar = 10 μm in (J).

### Comparison of mRNA expression levels of *NPGL* and *NPGM* between male and female

At six weeks of age, the body mass of females was significantly larger than that of males, while blood glucose levels remained unchanged (Fig. 4A, B). Real-time PCR showed that the *NPGL* mRNA expression level in females was significantly higher than that in males (Fig. 4C). In contrast, the *NPGM* mRNA expression level did not change between males and females (Fig. 4C). Among other feeding-related factors, the *GAL* mRNA expression level in females tended to be higher than that in males (Fig. 4C).

**Fig. 4.**
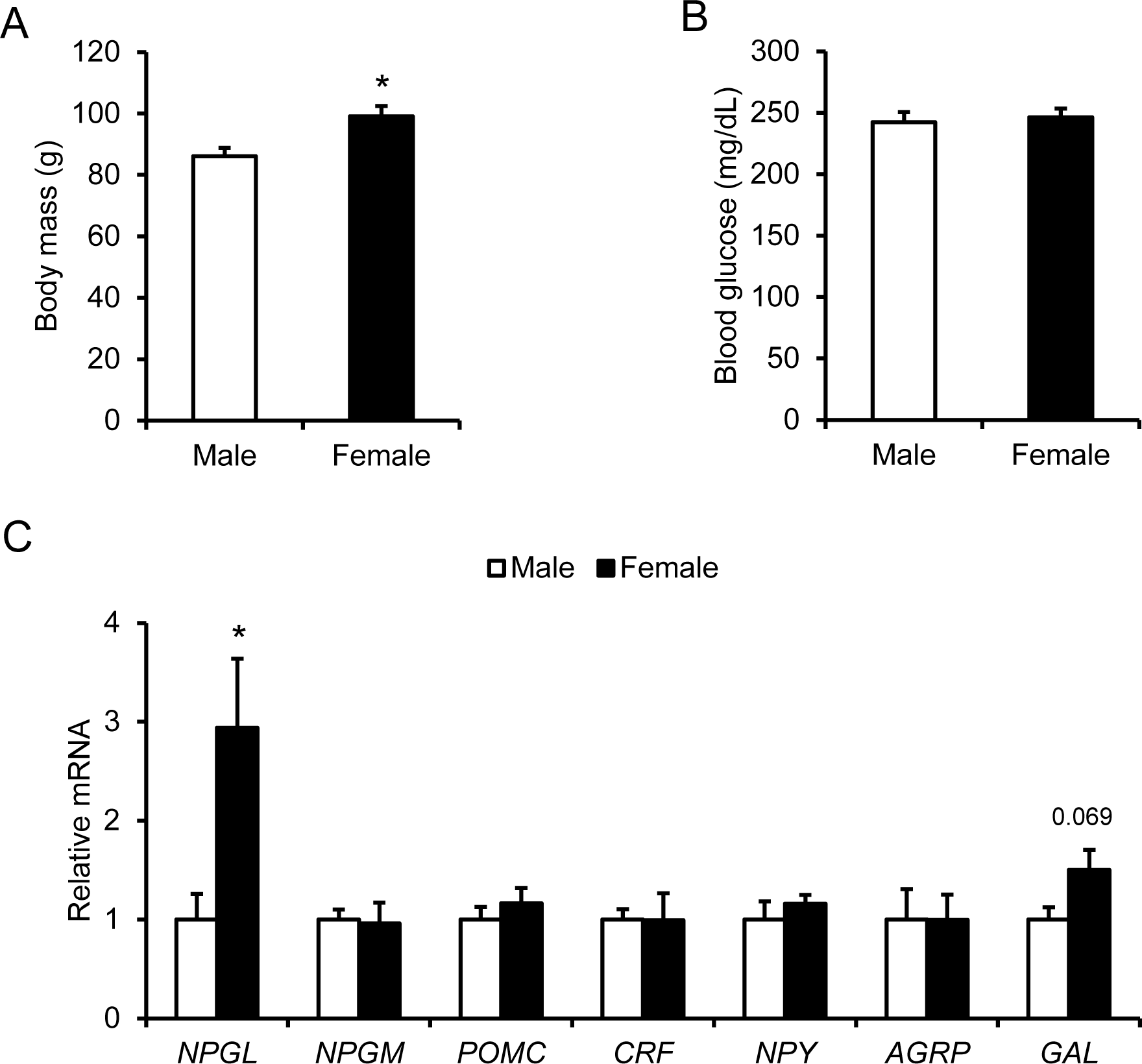
Comparison of body mass (A), blood glucose levels (B), and gene expression of hypothalamic factors (C) between males and females are shown. Data are expressed as the mean ± SEM (*n* = 5). Data were analyzed by Student’s *t*-test. An asterisk indicates a statistically significant difference (\**p* < 0.05).

*POMC*, *CRF*, *NPY*, and *AGRP* mRNA expression levels did not differ between males and females (Fig. 4C).

### Response to the food deprivation for 24 hours

After 24 h of food deprivation, males showed a trend toward decreased body mass and significantly decreased blood glucose levels (Fig. 5A, B). Females showed significant decreases in body mass and blood glucose levels (Fig. 5A, B). Real-time PCR showed that *NPGL*, *NPGM*, *NPY*, and *AGRP* mRNA expression levels were significantly increased after 24 h of fasting in male quails (Fig. 5C). In female quails, *NPGM*, *NPY*, and *AGRP* mRNA expression levels were significantly increased, whereas *NPGL* mRNA was not changed after 24 h fasting (Fig. 5D).

**Fig. 5.**
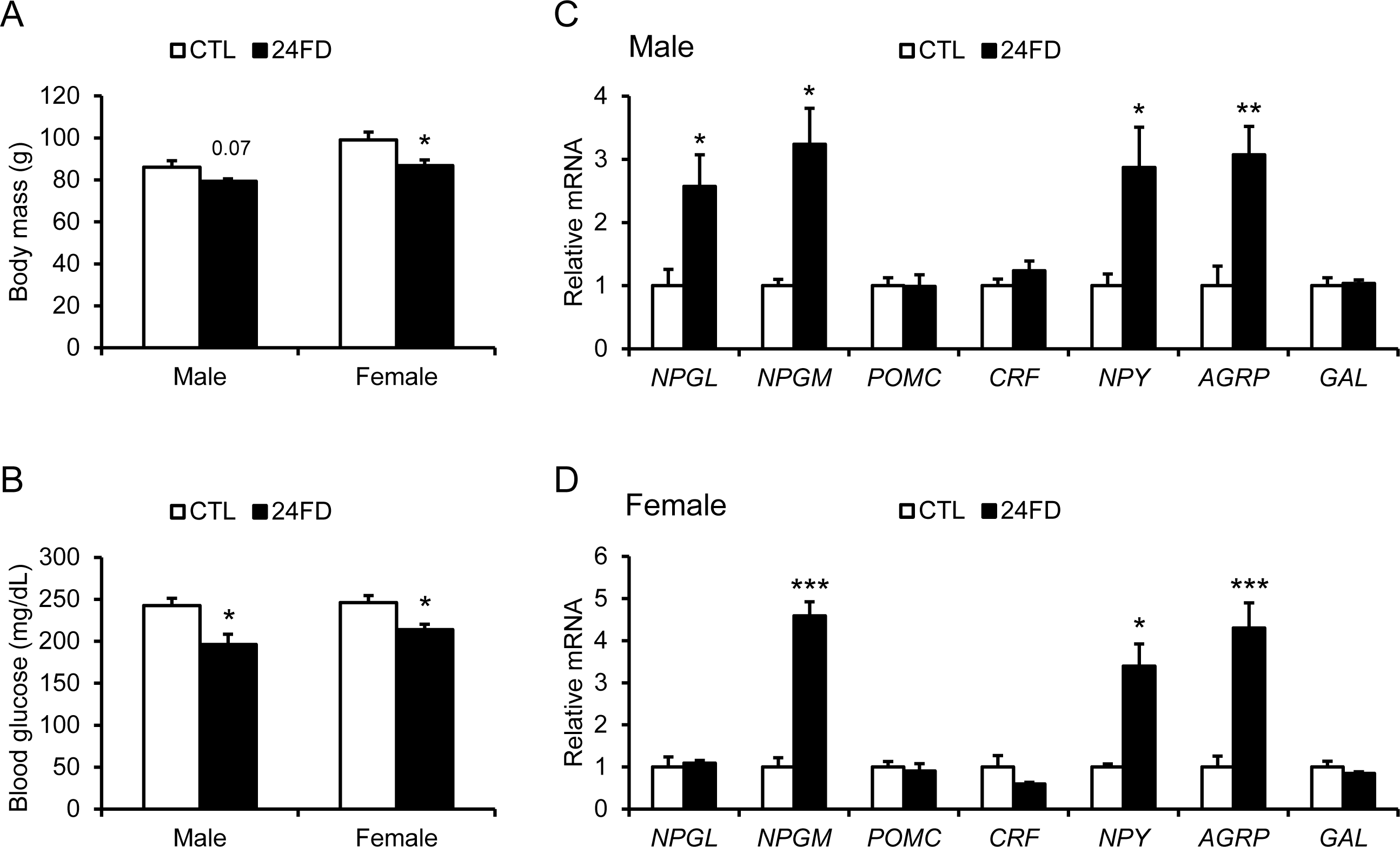
Analysis of the effects of food deprivation for 24 h in males and females. (A) Comparison of body mass between control (CTL) and fasted (24FD) groups. (B) Comparison of blood glucose levels between control and fasted groups. (C) Comparison of mRNA expression levels of hypothalamic factors in male hypothalamus. (D) Comparison of mRNA expression levels of hypothalamic factors in female hypothalamus. Data are expressed as the mean ± SEM (*n* = 5). Data were analyzed by Student’s *t*-test. Asterisks indicate a statistically significant difference (\**p* < 0.01, \*\**p* < 0.01, \*\*\**p* < 0.005).

## DISCUSSION

The hypothalamus is one of the centers of feeding and energy metabolism (Huang et al., 2021). These mechanisms are regulated by neuropeptides and/or hormones (Schwartz et al., 2000). NPGL and NPGM have been reported as novel neuropeptides related to energy metabolism in rodents and chickens (Iwakoshi-Ukena et al., 2017; Shikano et al., 2019; Kato et al., 2021; Narimatsu et al., 2022a, b). NPGL and NPGM have no similarity to other neuropeptides. In this study, we identified cDNAs encoding precursor proteins of NPGL and NPGM to investigate the biological characterization of NPGL and NPGM in Japanese quail. Moreover, we elucidated the localization of mRNA-expressing cells and immunoreactive cells in the hypothalamus, mRNA expression levels in sex differences, and the effect of food deprivation on these expressions.

Comparing the precursor proteins of NPGL and NPGM, the apparent difference is that the total lengths (181-185 amino acids) of NPGL are longer than those (132-139 amino acids) of NPGM in mammals and birds (Fig. 1). This indicates that the *NPGL* and *NPGM* genes were paralogized by gene duplication and conserved among each gene in higher vertebrates. The similarities in amino acid residues between quails and chickens were 91% in the precursor protein of NPGL, 99% in the mature protein of NPGL, 92% in the precursor protein of NPGM, and 100% in the mature protein of NPGM. These results suggest that *NPGL* and *NPGM* genes are almost maintained during the process of divergence between the same galliform species, quails, and chickens. The mature NPGL of quails showed 84% and 99% identity with those of mammals (humans, rats, and mice) and chickens, respectively (Fig. 1). In addition, the position of cysteine residues in the mature protein was consistent with previous reports (Shikano et al., 2018a; Ukena, 2018; Huang et al., 2022). Therefore, it is likely that the tertiary structure of mature NPGL is similar in quails and other vertebrates. In contrast, the mature NPGM of quails showed 54%, 52%, and 100% identity with those of humans, rodents (rats and mice), and chickens, respectively. Moreover, although mature NPGM in quails and chickens contains two cysteine residues, mature NPGM in mammals contains three cysteine residues. Thus, the tertiary structure of NPGM in mammals may differ from that of birds. Given that the findings of the tertiary structure provide convincing evidence for the physiological role of neuropeptides (van Vlijmen et al., 2004), these results imply that the structural features and physiological roles of NPGM diversify during evolution from birds to mammals. Further studies on the relationship between the three-dimensional structures of NPGM and their physiological functions in vertebrates are required.

Quantitative PCR analysis revealed that the mRNA expression of *NPGL* was upregulated by food deprivation in male quails. Food deprivation also upregulated the mRNA expression of potent orexigenic factors such as *NPY* and *AGRP* mRNAs in quails in the present and previous studies (Boswell et al., 2002). In situ hybridization revealed that some of the *NPGL* mRNA-expressing cells were localized to the IN in male quail. The IN in the hypothalamus is the center of feeding behavior in quails, where NPY and AGRP are also expressed (Boswell et al., 2002). Furthermore, our previous studies have shown that NPGL promotes feeding behavior in male chicks and mice (Matsuura et al., 2017; Shikano et al., 2018b, 2019). These data suggest that NPGL might have an orexigenic effect in male quails.

In contrast to males, we observed steady expression of *NPGL* under food deprivation in females, despite the higher expression in females than in males. These results suggest that endogenous NPGL may work not only in energy deprivation but also in other conditions in females. Regarding this sex difference, it is possible that NPGL is involved in growth, fat accumulation, and lipid production in female quails. At the onset of sexual maturity, weight and blood triglyceride levels increase progressively due to the acceleration of lipid metabolism to obtain the lipids needed for egg production in female quails (Furuse et al., 1991; Baer et al., 2015). Indeed, previous studies in chickens and rodents have shown that NPGL promotes weight gain due to fat accumulation in adipose tissue and the liver (Iwakoshi-Ukena et al., 2017; Shikano et al., 2019; Narimatsu et al., 2022a). Moreover, we found that *GAL* mRNA expression tended to increase in females. Galanin is a well-known orexigenic factor in birds and mammals (Leibowitz, 2007; Tachibana et al., 2008). Morphological analysis revealed that parts of the NPGL-immunoreactive cells were co-localized with galanin- immunoreactive cells in mice (Shikano et al., 2020). Therefore, these data imply that lipid metabolism during sexual maturation in female quails may be regulated by NPGL and galanin. We need to investigate the effects of NPGL and galanin on lipid metabolism in adipose tissue and the liver during female sexual maturation, especially egg production, in detail.

The PCR analysis also showed equivalent mRNA expression of *NPGM* between males and females. Considering that the mRNA expression of *NPGM* was upregulated by fasting in both males and females, these results suggest that NPGM may act as an appetite regulator, regardless of sex. Morphological analyses showed that *NPGM* mRNA and NPGM-immunoreactive cells were localized in the MM of the hypothalamus. Since NPGM is also expressed in the MM of chickens and is upregulated by fasting (Shikano et al., 2018a; Kato et al., 2022), the MM may play a key role in the physiological function of NPGM among birds. We previously reported that chicken NPGM was produced in histaminergic neurons of the MM, and i.c.v. administration of histamine or NPGM suppressed feeding behavior (Bessho et al., 2014; Shikano et al., 2018a). Therefore, histaminergic neurons containing NPGM in the MM appear to be involved in maintaining basal levels of food intake in birds. In addition to the cell bodies, we observed NPGM-immunoreactive fibers projected to the external layer of the ME in the quail. Several reports have shown that neurons projected to the external layer of the ME often organize neural circuits with fenestrated capillaries and secrete neuropeptides or neurotransmitters as hormones (Yamada and Mikami, 1985; Walsh and Kuenzel, 1997; Tsutsui et al., 2000; Ezzat et al., 2006). These findings indicate that NPGM acts as a neurohypophysial hormone-regulating peptide in quails. Future studies are necessary to unveil the physiological role of NPGM neurons located in the MM or projected to the external layer of the ME in quails.

Although *NPGL* mRNA-expressing cells were localized in the IN and ME, *NPGM* mRNA-expressing cells and NPGM-immunoreactive cells were only localized in the MM. In chickens, both NPGL and NPGM neurons are localized in the IN and the MM (Shikano et al., 2018a). The quail in this study was used after sexual maturity, whereas the chicken used in the previous study was a juvenile. Thus, it is possible that the expression regions of NPGL and NPGM change greatly before and after sexual maturity in birds. In future studies, we will analyze the localization of NPGL and NPGM in juvenile quails. In addition, previous studies have reported that perikarya, which expresses neuropeptides related to energy metabolism, is detected in the ME. NPY is produced in the ME of chickens and golden hamsters (Sabatino et al., 1987; Walsh and Kuenzel, 1997), whereas POMC is expressed in the ME of rats (Wittmann et al., 2017). In this study, the cell size of *NPGL* mRNA-containing cells in the ME was very small. These cells in the ME may be tanycytes, which are epithelial cells.

Tanycytes play an important role in modulating energy balance because they regulate the secretion of neurohormones from the hypothalamus to the hypophysial portal vein via the cerebral spinal fluid (Dali et al., 2023). These data suggest that the ME may act not only as a terminus connected with the pituitary gland in neurosecretion, but also as a center in energy metabolism. Further studies are required to evaluate the regional changes in NPGL/NPGM-expressing neurons associated with growth and the effects of NPGL expressed in the ME on energy metabolism.

In the present study, NPGL-like immunoreactive cells were not detected. In mice, colchicine treatment, which inhibits intracellular axonal transport, is used to detect NPGL-immunoreactive cells (Matsuura et al., 2017). Given that the lack of NPGL-immunoreactive signals was caused by the absence of colchicine treatment in this study, these data support that NPGL is constantly secreted and does not accumulate intracellularly in quails and rodents. In contrast to NPGL, colchicine treatment is not necessary to detect NPGM in cell bodies. We speculate that NPGM is stored intracellularly and may be secreted upon specific signaling inputs. Additional studies focusing on the secretory mechanisms of NPGL and NPGM are required to unravel their mechanisms of action.

In conclusion, we unmasked the similarities or differences between NPGL and NPGM in Japanese quail from the viewpoint of molecular biology and histology. We identified *NPGL* and *NPGM* cDNAs encoding precursor and mature proteins from the hypothalamus of Japanese quail and characterized the localization of mRNA-expressing cells and immunoreactive cells, respectively. The mRNA expression of *NPGL* is affected by sex, suggesting that endogenous NPGL may have different roles in males and females. In contrast to NPGL, the mRNA expression level of *NPGM* was independent of sex. In addition, *NPGM* mRNA expression was upregulated by fasting. This is the first report of NPGL and NPGM in the Japanese quail. Judging from the abundant advantages in biological research using quails, this study will provide important clues to the physiological functions of NPGL and NPGM in the energy metabolism of birds. In future studies, we will perform acute or chronic i.c.v. administration of NPGL and NPGM in both male and female quails to reveal the physiological function during sexual maturity.

## ACKNOWLEDGMENTS

We are grateful to Mr. Shogo Moriwaki for experimental support. This work was supported by KAKENHI Grants (JP20KK0161 and JP22H00503 to KU, JP16K07440 to EI-U, and JP22KJ2331 to MK) and the Hiroshima University Graduate School Research Fellowship (MK and YN).

## AUTHOR CONTRIBUTIONS

MK and KU designed the experiments, and MK, EI-U, YN, and MF performed the experiments and analyzed the results. MK wrote the first draft of the manuscript.

MK, YN, and KU reviewed and edited the manuscript. All authors have read and agreed to the published version of the manuscript.

